# Molecular Graph Contrastive Learning with Parameterized Explainable Augmentations

**DOI:** 10.1101/2021.12.03.471150

**Authors:** Yingheng Wang, Yaosen Min, Erzhuo Shao, Ji Wu

## Abstract

Learning generalizable, transferable, and robust representations for molecule data has always been a challenge. The recent success of contrastive learning (CL) for self-supervised graph representation learning provides a novel perspective to learn molecule representations. The most prevailing graph CL framework is to maximize the agreement of representations in different augmented graph views. However, existing graph CL frameworks usually adopt stochastic augmentations or schemes according to pre-defined rules on the input graph to obtain different graph views in various scales (e.g. node, edge, and subgraph), which may destroy topological semantemes and domain prior in molecule data, leading to suboptimal performance. Therefore, designing parameterized, learnable, and explainable augmentation is quite necessary for molecular graph contrastive learning. A well-designed parameterized augmentation scheme can preserve chemically meaningful structural information and intrinsically essential attributes for molecule graphs, which helps to learn representations that are insensitive to perturbation on unimportant atoms and bonds. In this paper, we propose a novel Molecular Graph Contrastive Learning with Parameterized Explainable Augmentations, MolCLE for brevity, that self-adaptively incorporates chemically significative information from both topological and semantic aspects of molecular graphs. Specifically, we apply deep neural networks to parameterize the augmentation process for both the molecular graph topology and atom attributes, to highlight contributive molecular substructures and recognize underlying chemical semantemes. Comprehensive experiments on a variety of real-world datasets demonstrate that our proposed method consistently outperforms compared baselines, which verifies the effectiveness of the proposed framework. Detailedly, our self-supervised MolCLE model surpasses many supervised counterparts, and meanwhile only uses hundreds of thousands of parameters to achieve comparative results against the state-of-the-art baseline, which has tens of millions of parameters. We also provide detailed case studies to validate the explainability of augmented graph views.

**CCS CONCEPTS:** • Mathematics of computing → Graph algorithms; • Applied computing → Bioinformatics; • Computing methodologies → Neural networks; Unsupervised learning.

## 1 INTRODUCTION

The capability of obtaining informative molecular representations is an essential key and a crucial prerequisite in AI-driven drug design and discovery. Well-designed molecular representations will benefit various academic areas and industrial applications such as improving rational chemical design, reducing R&D cost, decreasing the failure rate in potential drug screening trials, as well as speeding the process of *de novo* drug design [65]. However, due to the large-scale vocabulary of possible stable chemical compounds, it is challenging to develop generalizable, transferable, and robust molecular representations covering the entire molecule space. Traditional molecular fingerprints like SMILES [58] and ECFP [46], provide a description of a specific part of the molecular structure [13]. However, they usually require intensive manual feature engineering and strong domain knowledge, and finally become highly task-dependent, with no generalizability for other new-come tasks.

Besides, these methods also suffer two main problems. Firstly, **most of them are task-specific**. For many areas like computer vision (CV) [20] and speech signal processing (SSP) [15], the number of labeled samples could easily reach several million or even more. However, it is not the same situation with molecule-related tasks, which often require an expensive and time-consuming process to get access to labeled molecule data. The scenario with unlimited unlabelled data while a tiny portion labeled ones are similar to natural language modeling. Thus, the state-of-the-art natural language pre-training techniques [6, 7] can be easily applied over string-based molecule fingerprints. But, these approaches will lead to the second problem: **string-based representations fail to encode important topology**. Molecules can be naturally represented by molecular graphs which preserve rich structural information. Over the past few years, graph representation learning has emerged as a powerful strategy for analyzing graph-structured data, which has also been extended to encode molecular graphs. Among them, the most prevailing graph neural networks (GNN) [8, 27, 29] receive considerable attention, which aims to transform graph components to low-dimensional dense embeddings that preserve attributive and structural features. Despite existing GNN models flourish in many tasks, such as molecular property prediction [56] and virtual screening [57], they are mostly established in a supervised manner [44], bringing back to the first problem.

Recently, Contrastive Learning (CL), as the revitalization of the classical information Maximization (InfoMax) principle [32], has performed great achievement in many fields, e.g., visual representation learning [4, 19] and natural language processing (NLP) [38]. The CL framework seeks to maximize the mutual information (MI) between the input (i.e. images) and its representations (i.e. image embeddings) by contrasting positive pairs with negative-sampled counterparts. Inspired by the previous success of the most famous Deep InfoMax (DIM) [2], Deep Graph InfoMax (DGI) [55] firstly introduces the InfoMax principle into GNN-based self-supervised learning. DGI simply applies random shuffling on node features to generate graph augmentations. Then, a contrastive objective is used to maximize the MI between node patch embeddings and graph summary representations. Following DGI, GRACE [72] generates corrupted views of graphs by applying random node feature masking and edge removing and measures MI from two views both in patch level. GraphCL [69] proposes four simple graph augmentations including node dropping, edge perturbation, attribute masking, and induced subgraph. To supplement more global information, MVGRL [18] firstly utilizes graph diffusion kernels [31] to augment graphs and constructs graph views by randomly sampling subgraphs. Then it contrasts patch and graph representations across views. Despite these methods achieve success in both node- and graph-level tasks, all of them only perform simple data augmentations but ignore exploring more abundant ones, which proved to be a critical component for visual representation learning [59]. GCA [73] studies the discrepancy in the impact of nodes and edges when performing data augmentation in graph CL. It identifies important edges and feature dimensions via centrality measures and then generates different graph views by masking the unimportant ones.

Although graph CL frameworks with simple or adaptive graph augmentations have shown significant improvements on the social network, citation network, and other internet datasets, they can not be easily transferred to biomedical molecule datasets because such augmentations will **generate meaningless graph views at the molecule-level**. Uniformly dropping nodes or perturbing edges may drastically change the identity and even the validity of a compound, e.g., removing an aromatic bond or dropping an atom from a functional group. Besides, centrality-based adaptive graph augmentations largely borrow the hypothesis from network science that centrality reflects inter-node influences. Thus, the generated graph views are constructed by the most influential nodes, which will also deteriorate meaningful substructure in molecular graphs. Meanwhile, from the attribute perspective, the simple augmentations maybe **not sufficient to generate diverse contexts for nodes** and **improperly mask important feature dimensions**, e.g., masking feature dimensions like inter-atomic distances when predicting targets related to atomic orbital and energy gap. Therefore, such operations will cause pointless augmentations, leading to difficulty in optimizing the contrastive objective.

To this end, we propose a novel contrastive framework for selfsupervised molecular graph representation learning, as shown in Figure 1, which we refer to as Molecular Graph Contrastive Learning with Parameterized Explainable Augmentations, MolCLE for brevity. (i) In MolCLE, we first design two parameterized graph data augmentations on both topology and attribute level. We apply a generative probabilistic model to learn *intrinsic underlying molecular graph structures* as the topology-level augmentations, which are believed to make the most contribution to molecule representations readout from GNNs. Simultaneously, MolCLE also learns *feature selectors* that mask out unimportant atom features to generate attribute-level augmentations. (ii) Then, we utilize them to obtain correlated molecular graph views and train the model using a contrastive loss to maximize the agreement between augmented molecular graph representations in these two views. (iii) We pretrain MolCLE with millions of unlabelled molecule data, and then leverage the pre-trained GNN backbones to many downstream molecular property prediction datasets followed by task-specific fine-tuning. With only less than a million parameters, our approach achieves better performance than state-of-the-art methods with hundreds of million parameters. Extensive experiments show that MolCLE outperforms existing methods and our self-supervised method even surpasses its supervised counterparts on many downstream tasks.

**Figure 1:**
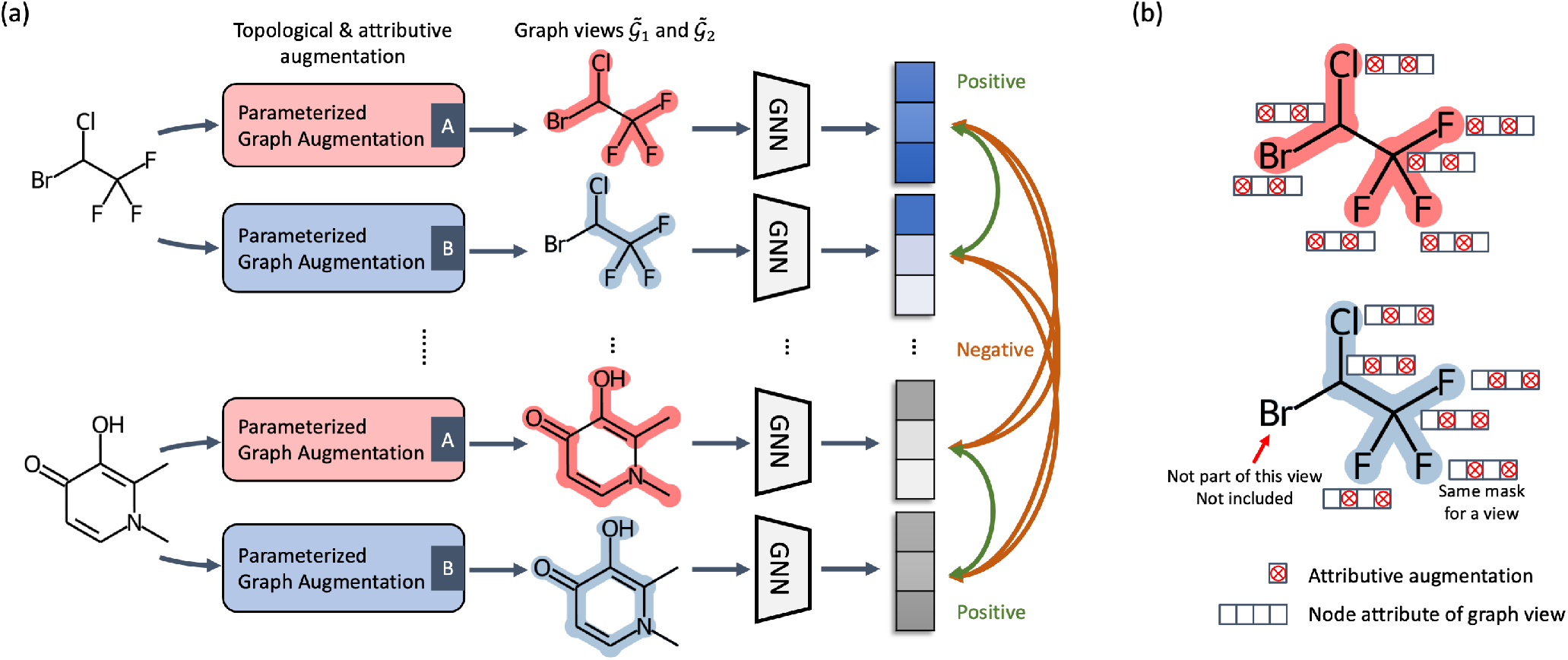
Our proposed <u>Mol</u>ecular Graph <u>C</u>ontrastive <u>L</u>earning with Parameterized <u>E</u>xplainable Augmentations (MolCLE) model. (a) Molecular contrastive learning pipeline. We apply two parameterized graph augmentation components for each molecule, where they simultaneously learn topological and attributive augmentation while retaining domain priors, to generate two explainable augmented graph views. Finally, the augmented graph views from the same molecule are treated as positive pairs, pulling their representations closer in the following contrastive training process. And the negative samples are constructed from the anchor and augmented views of other graphs in the same minibatch. (b) Parameterized augmentation illustration. For each graph view, certain topological connectivity and attribute dimensions are selected, where atoms in the same view share a common attribute selector. Noted that some atoms’ attributes will not be included if they are ignored by the topology selector.

Our contributions of this paper could be summarize as:

Firstly, we propose a contrastive learning framework for selfsupervised molecular graph representation learning with parameterized, explainable data augmentation. The proposed MolCLE framework jointly performs data augmentation on both topology and attribute levels that can automatically emphasize important parts of the graph structure and attributes, which encourages the model to extract essential features from both aspects.

Secondly, we conduct comprehensive empirical studies using 6 downstream popular benchmark datasets on molecular property prediction compared with many current state-of-the-art results for each dataset. We observe a huge relative improvement of MolCLE over existing baselines that even surpasses some supervised counterparts.

## 2 RELATED WORK

### 2.1 Molecule Representation Learning

To represent molecules in the embedding space, traditional chemical molecule fingerprints, such as Morgan Identifier [39] and ECFP [46], try to encode the neighboring atoms of the central atom in the molecule into a fix-length vector. To improve the expressive power of chemical fingerprints, some studies [8] introduce learning-based methods like convolutional layers to learn the neural fingerprints of molecules. Then the learned molecule representations are applied to downstream tasks. Following these works, many other researchers take the SMILES representation [58] as input and use language modeling approaches to extract molecule fingerprints like Mol2vec [24] and Seq2seq [65]. Recently, considering molecule naturally as graph-structure data, many works [27, 56] explore the flourished GNNs to encode molecular graphs into low-dimensional representations. A couple of works [54, 71] extend attention mechanism to graph aggregation operation by learning the aggregation weights automatically. Moreover, Gilmer et al. [12] proposes a unified message passing framework to encode molecular graphs for quantum chemical property prediction. Following this work, some research works [34, 66] propose to model bond interactions over this framework. Besides, several works [34, 57] model molecular graphs and cross-molecule interactions in a multi-level manner. More recently, several self-supervised GNN-based models [23, 47] tailored for molecule data design elaborate pre-training techniques and can obtain good molecule representations, which achieve great performance on downstream tasks.

### 2.2 Graph Neural Networks

Recent success of GNNs [8, 12, 29] have aroused many attention to use these methods for analyzing various graph-structured data. The general framework includes an iterative neighboring aggregation (namely message passing) scheme to update nodes’ hidden states within information from their neighboring nodes and themselves. Let 𝒢 ={𝒱, ε} denote an undirected graph, ***h***_*n*_ is the embedding of the vertex *v*_*n*_ ∈ 𝒱. Considering a *K*-layer GNN *f*_*GN N*_(·), the propagation of the *k*th layer if represented as:

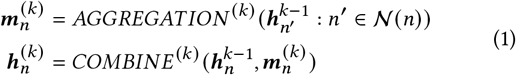

where 𝒩 (*n*) is a set of neighboring vertices adjacent to *v*_*n*_, and *AGGREGAT ION* ^(*k*)^ (·) and *COMBI N E* ^(*k*)^ (·) are component functions of the GNN layer, which have multiple popular choices like mean, max pooling and attention mechanism [53]. After the K-layer propagation, the graph-level embedding for 𝒢 is summarized on layer embeddings through the READOUT function:

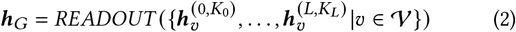

### 2.3 Contrastive Representation Learning

The main idea of contrastive learning is aiming to make representations agree with each other under proper transformations, raising a recent surge of interest in visual representation learning [4, 19]. For visual data, negative samples can be easily generated using a multistage augmentation pipeline [2, 10] including rotation, resizing, cropping, flipping, color jittering and distortion, etc. Existing works [52, 59] propose to store negative samples with a memory bank. In contrast, some other works [4, 67] explore in-batch negative samples. Considering an image patch as the anchor, the positive samples are often constructed by patches in neighboring views [21] or a global summary vector [22]. Then, the positives are contrasted with the negative counterparts like patches of other images [22].

On a parallel note, for graph data, traditional methods [30] trying to reconstruct the adjacency information of nodes [17], which could be seen as a contrast of locality. These methods emphasize the proximity information over the topological information [45], which ignores information from attributes. Also, these methods are known to be error-prone with inappropriate hyperparameter tuning [16, 43]. The most famous DGI [55] is the first work that combines GNN and CL, which applies an InfoMax objective and maximizes MI between global graph-level and local node-level embeddings. Following DGI, GMI [42] measures MI between input and representations of both nodes and edges without data augmentations by two discriminators. GRACE [72] focuses on maximizing the agreement of node embeddings across two corrupted views of the graph. GraphCL [69] proposes four graph data augmentations to generate different views and maximizing the agreement of these views. MVGRL [18] performs graph diffusion kernels to learn both node and graph representations and then contrast them. InfoGraph [51] applies a discriminator with a batch of graphs to determine which patches are from the same graph. GCA [73] introduces two adaptive graph data augmentations, aiming to discover the centrality-based influential substructures of the graph. However, these methods do not explicitly consider chemical and biological priors inside molecular graphs, leading to improper graph augmentations at both structural and attribute levels. Also, these augmented views will deteriorate optimization of the contrastive objective and reach a suboptimal performance.

#### Connection to related works

We summarize the comparisons between our proposed MolCLE method with other related state-of-the-art graph contrastive learning models, including DGI [55], GRACE [72], GraphCL [69], GMI [42], InfoGraph [51], MVGRL [18], and GCA [73] in Table 1. It is obvious that only our MolCLE model proposes parameterized explainable graph data augmentations on both topology and attribute levels for preserving domain priors better, which are especially crucial on molecular graph data.

**Table 1:** Summary of comparisons.

| Model | Augmentation |  |  | Contrastive objective |  |
| --- | --- | --- | --- | --- | --- |
|  | Topo. | Attr. | Type |  |  |
| DGI [55] | ✓ | ✗ | Uniform | Node | Graph |
| GRACE [72] | ✓ | ✓ | Uniform | Node | Node |
| GraphCL [69] | ✓ | ✓ | Uniform | Graph | Graph |
| GMI [42] | ✗ | ✗ | None | Node | Node |
| InfoGraph [51] | ✗ | ✗ | None | Node | Graph |
| MVGRL [18] | ✓ | ✗ | Uniform | Node | Graph |
| GCA [73] | ✓ | ✓ | Adaptive | Node | Node |
| **MolCLE** | ✓ | ✓ | Self-adaptive | Graph | Graph |

## 3 METHODOLOGY

In the following section, we introduce our proposed MolCLE model in detail, starting with the parameterized graph data augmentations, followed by the overall graph contrastive learning framework.

### 3.1 Preliminaries

Let 𝒢 = (𝒱, ε) represent the graph with 𝒱= {*v*_1_, *v*_2_, …, *v*_*N*_} denoting the node set and ε ∈ 𝒱 × 𝒱as the edge set. The numbers of nodes and edges are denoted by *N* and *Mi*, respectively. A graph can be described by an adjacency matrix ***A*** ∈ {0, 1} ^*N* ×*N*^, with *a*_*i j*_ = 1 if there is an edge connecting node *i* and *j*, and *a*_*i j*_ = 0 otherwise. Nodes in 𝒱 are associated with the *d*-dimensional features, denoted by ***X*** ∈ ℝ ^N×*d*^. As stated in Section 2.2, the message and hidden state representation of node *v*_*n*_ in the *k*th layer denotes 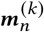 and 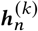, respectively. And the hidden representation of the last GNN layer serves as the final node representation 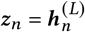, which can be then readout to a graph representation ***h***_*G*_.

### 3.2 Parameterized Graph Augmentation

The core idea of CL is maximizing agreement between augmented views to obtain *perturbation-invariant* representations [62]. Existing graph data augmentation mainly lies in two categories: generative-based methods and simple random disturbance. The former often requires prohibitive additional computation cost [1] and the latter usually destroy domain priors that deteriorate the quality of the learned representations [69]. Thus, it is obvious that self-adaptive graph data augmentation preserving underlying essential graph structures still remains under-explored. In our MolCLE model, we propose to design parameterized explainable augmentation schemes that self-adaptively learn to focus on intrinsically important graph topology and node attributes while perturbing relatively unimportant edges and features. Specifically, MolCLE learns *graph topology masks* and *feature selectors* using generative probabilistic models which select the important subgraph and mask out unimportant node features of input molecular graph as topology- and attribute-level augmentations, respectively. We argue that augmented graph views in molecule modeling scenarios should preserve chemically meaningful topological information instead of randomly dropping edges. Also, target-influential attributes should be emphasized to preserve semantic patterns rather than randomly masking out them.

#### 3.2.1 Topological augmentation

Previous topological augmentation methods usually apply uniform node dropping, edge perturbation [69] or graph corruption guided by rules in specific domains [73]. However, these approaches can not be adapted to molecular graph augmentation. They may generate meaningless augmentation with no chemical significance. Thus, we consider a self-adaptive approach that automatically learning intrinsic underlying topology for input molecular graphs. Inspired by previous works on explaining GNNs [35, 68, 70], we utilize a generative probabilistic model to obtain topological augmentations. Different from these works, we do not have training labels in contrastive learning, There-fore, we try to find a graph view *G*_*A*_ by maximizing the MI between graph embeddings obtained from original graph topology *G*_*O*_ and the underlying graph topology *G*_*A*_:

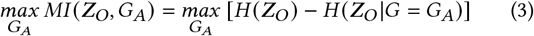

where ***Z***_*O*_ is the graph embeddings output from the GNN model with *G*_*O*_ as the input. The MI measures the proximity of the graph embedding ***Z***_*A*_ to ***Z***_*O*_ when the input graph to the GNN model is limited to the augmented graph view. Maximizing MI can help to find the underlying graph view that makes the most contributions to generate ***Z***_*A*_ closest to ***Z***_*O*_. The intuition behind this is that the remaining topology of the graph view *G*_*A*_ should preserve the original graph embedding ***Z***_*O*_ as much as possible, e.g., if removing an edge dramatically disturbs ***Z***_*O*_, thus, the edge is essential and should be selected in *G*_*A*_. Otherwise, it can be removed for its irrelevance.

Due to the intractable number of potential subgraphs hindering the model from optimizing the objective directly [35], we follow Janson et al. [26] to consider a direct way that factorizing the probability of a specific augmented graph view:

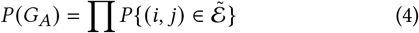

where 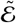 denotes the edge set of the graph view, and 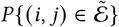 is drawn from a Bernoulli distribution 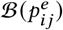. Considering 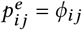 and *G*_*A*_ is drawn from the distribution *d* ^(Φ)^ of the *ϕ*-parameterized graph view, we reformulate Equation (3) as:

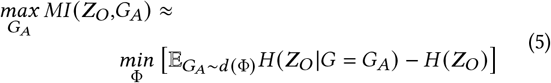

Besides, for efficiently optimizing Equation (5) with gradient descent [68], we also adopt the reparameterization trick [25] to relax edge sampling weight variables 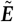 from binary to continuous between 0 to 1. Formally, the entry of 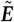 can be instantiated as a binary concrete distribution [36]:

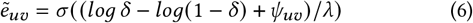

where *δ* is drawn from a Uniform distribution 𝒰 (0, 1) and *σ* (·) denotes the Sigmoid function. We parameterize *ψ*_*uv*_ ∈ R by a fullyconnected neural network *FCNN*_Γ ([_***z***_*u*_ ; ***z***_*v*])_, where [;] represents a concatenation operation.

Although Maddison et al. [36] have demonstrated that using binary concrete distribution to approximate the Bernoulli distribution is reasonable, the conditional entropy is still remained difficult to be optimized. Inspired by Ying et al. [68], we modify it with a soft cross-entropy 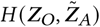 and adopt Monte Carlo [37] to complete approximate optimization for Equation (5):

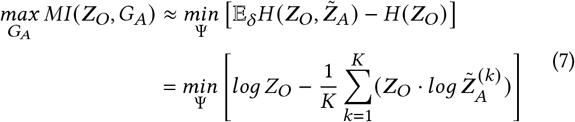

where 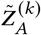 denotes the *k*th sampled graph view with Equation (6) and *K* is the total number. In practice, an *m*-dimensional Softmax layer is applied upon ***Z***_*O*_ and 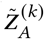 for stable optimization. Thus, Equation (7) can be rewritten as follows:

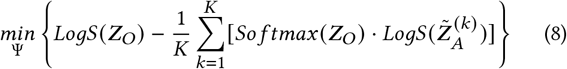

where *LogS* (·) combines the logarithm with the *Softmax* (·) function. The intuition behind is obvious that the graph embedding with a well-augmented graph view as GNN’s input should be closest to the original one.

Through optimizing Equation (8), the proposed model can easily learn appropriate topological augmentations for molecular graph instances, which include meaningful semantic information over molecular structures. We summarize the topological augmentation pipeline in Figure 2. However, the obtained graph views maybe not concerning constraints avoiding chemical meaninglessness. Therefore, we propose two regularization terms to preserve desired chemical properties.

**Figure 2:**
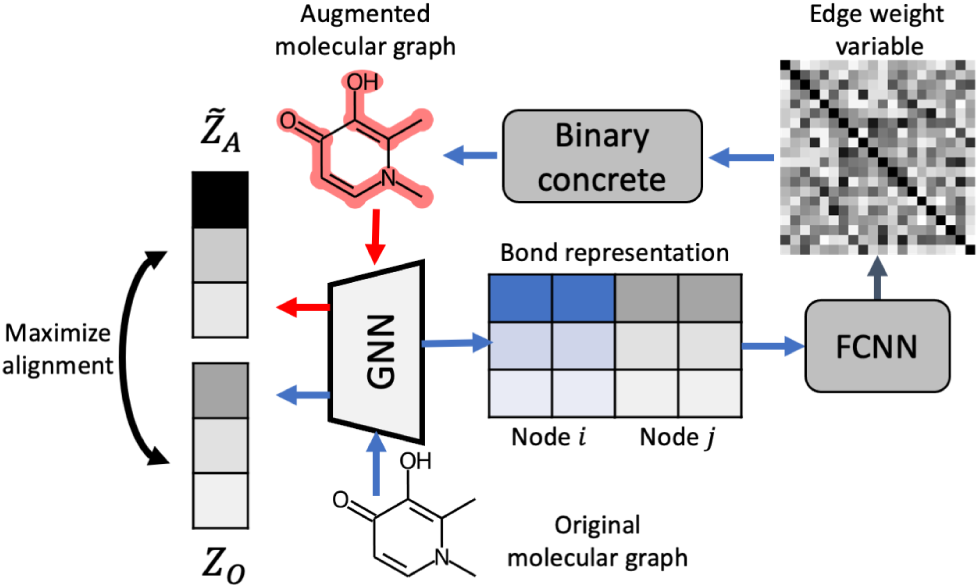
Topological augmentation.

#### Ratio constraint

Ying et al. [68] introduces a *size constraint* on the obtained explanatory graphs by adding || Φ ||_1_, the *l*_1_ norm on latent variables Φ, as a regularization term. But in our case, we need to preserve a reasonable size for augmented graph views instead of only penalizing the sum of all elements of Φ. Previous work [35] imposes an alternative solution, namely *budget constraint*, which applies a hard threshold *B* to limit the size of explanatory graphs.

However, it can not be adapted to molecular graphs having various sizes. Thus, we propose a *ratio constraint* that self-adaptively restricts the size of each molecular graph, i.e., ensuring the topological augmentation ratio. The ratio constraint can be formulated as follows:

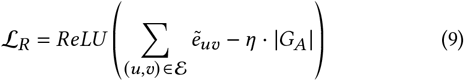

where *ReLU* (·) is the ReLU activation and *η* denotes the default topological augmentation ratio. The ratio constraint ℒ _*R*_ = 0 when the ratio of the size of the augmented graph view to the size of the original molecular graph is lower than the default ratio *η*.

#### Continuity constraint

In the molecular modeling scenario, the augmentation graph views are usually expected to be connected. Therefore, adding a constraint on generating continuous topological masks is necessary. The significance of this claim has been empirically proved and explicitly introduced [35, 68]. Thus, we design a more specific regularization term on the topological connectivity of augmented graph views:

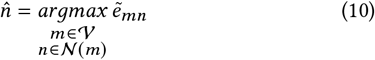

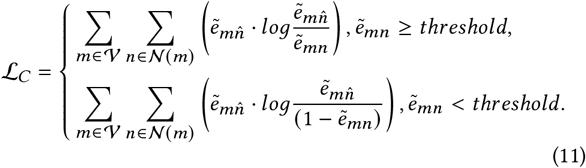

where 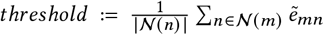. In practice, we add a uniform noise over 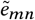 to avoid local optimum. Obviously, Equation (11) encourages the augmented graph views to include small connected subgraphs rather than many discretized edges.

#### Edge-entropy constraint

Although Equation (11) ensures the topological connectivity of augmented graph views, it will make the topological masks smoother, which means that different edge weight variables gradually become indistinguishable. Thus, we also apply the element-wise entropy [35, 68] constraint to increase the gap of edge variables 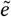 with large values and those with small values:

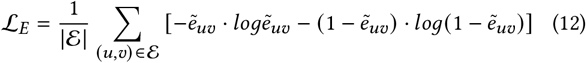

#### 3.2.2 Attributive augmentation

Existing attributive augmentation methods mainly concentrate on uniform or rules-guided attribute masking [69, 72, 73], ignoring to explore more strategies on self-adaptively selecting influential feature dimensions. Here we design learnable feature selectors to add noise on node attributes via automatically masking a fraction of dimensions with zeros in node features. Formally, MolCLE learns a binary feature selector 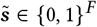 where each dimension 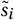 is drawn from a Bernoulli distribution 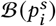. With ***x***_*A*_ representing all dimensions of node features in *G*_*A*_, the selected node features can be considered as the element-wise multiplication of the feature selector and ***x***_*A*_, which is formulated as 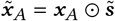. Thus, we rewrite Equation (3) to jointly optimize topological and attributive augmentation via maximizing the MI objective:

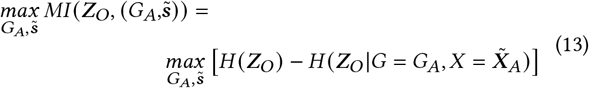

where 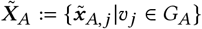 is obviously related to 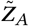 in Equation (7) and (8) but do not change the formulation of these two equations. Therefore, we still serve Equation (8) as the optimization objective. Here we provide some intuition behind this objective. If a specific feature is important, the feature selector will learn to select it, which means that the corresponding weights in 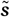 will be close to one, and vice versa.

However, similar to topological augmentation, for backpropagating gradients in Equation (13) to the feature selector 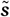 smoothly, we have to apply a reparameterization trick [25] on ***x***_*A*_:

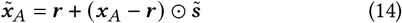

where ***r*** denotes a random variable drawn from the empirical distribution. Meanwhile, we also give some additional constraints for attributive augmentation, aiming to generate more meaningful graph views.

#### Size constraint

Similar to topological augmentation, we also need to keep a reasonable number of selected features in attributive augmentation. Thus, we propose a *size constraint* upon the number of selected feature dimensions like the setting in *ratio constraint*:

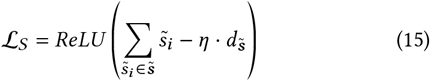

where *ReLU* (·)is the ReLU activation and *η* denotes the default attributive augmentation ratio. The ratio constraint ℒ _*S*_ = 0 when the ratio of the number of selected feature dimensions to the number of original feature dimensions is lower than the default ratio *η*.

#### Feature-entropy constraint

As we have done in Section 3.2.1, we also apply element-wise entropy to encourage the feature selector 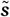 to be discrete:

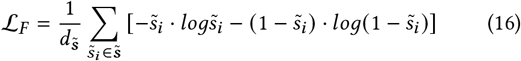

Finally, by jointly performing topological and attributive augmentation stated above, we can generate two augmented molecular graph views 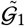 and 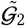. Considering these two augmentations as an overall augmentation pipeline, we perform it independently when generating each view but optimize the objectives in an end-to-end style with the contrastive objective in Section 3.3. Due to different initialization, proposed constraints, and the involved random variables, two generated views will display slightly different but chemically meaningful. We provide some visualized examples in Section 4.4 for more direct observation.

### 3.3 Graph Contrastive Learning Framework

We follow the general and widely-used graph contrastive learning framework [4, 22, 51, 69, 72, 73] which applies a contrastive objective to maximize the agreement of representations between different views. However, due to our graph-level learning scenario [51, 69], we employ the best-matched pipeline that enforces the encoded embeddings of each molecular graph in two different augmentation views to agree with each other and can be discriminated from embeddings of the others.

In our MolCLE model, at each iteration, we construct two augmentation pipelines 𝒜_1_ amd 𝒜_2_ and generate two correlated molecular graph views, denoted as 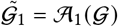 and 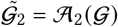, as a positive pair. Then, a GNN encoder *f*_*GN N*_ (·) defined in Section 2.2 will extract graph-level embeddings 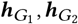 for augmented graphs 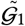 and 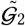. Following previous works [4, 69, 72, 73], we apply a projection head to map augmented graph embeddings to another latent space to calculate contrastive loss. The projection head is usually defined as a non-linear transformation, which can enhance the expressive power of the contrastive discrimination. Thus, in our study, we use a two-layer fully-connected neural networks to obtain representations 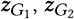 of 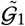 and 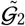 in the other latent space.

Then, we utilize a contrastive objective ℒ (·,·)to enforce maximizing the consistency between positive pairs 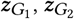 compared with negative pairs. For the *k*th graph *G*_*k*_ in a minibatch of *N* graphs, its augmented graph-level representation in latent space can be annotated as 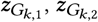 to form the positive sample. We serve the embedding in one view, i.e. 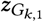 as the anchor. Naturally, the negative samples can be constructed from the anchor and augmented views of other graphs in the same minibatch [4, 5]. In our MolCLE model, we apply the wide-spread NT-Xent [40, 49, 61] loss as the contrastive objective and define the pairwise formulation over each positive sample:

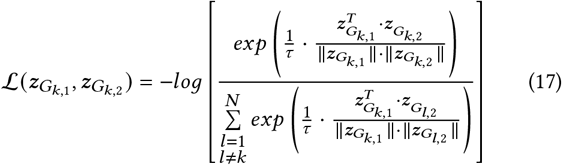

where *τ* is the temperature parameter. It should be noted that we compute the contrastive loss across all positive samples in a minibatch. The contrastive objective is jointly optimized with the molecular graph augmentation process. Equation (17) can be served as maximizing the MI between the representations of two augmented molecular graph views 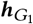 and 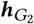, satisfying the classical InfoMax principle [32]. We follow Chen et al. [4] and You et al. [69] to allocate the same GNN encoder for two different views while there are many specific alternative CL frameworks [19, 22, 52].

## 4 EXPERIMENTS

In this section, we present the details of the experiments and the corresponding results. To illustrate the effectiveness of our proposed model, we compare it with some strong baselines on molecular property prediction tasks. We start with four research questions (RQs) to lead the experiments and the following discussions.

- **(RQ1)** Does our proposed MolCLE model outperforms existing baseline methods in molecular property prediction tasks over various datasets?
- **(RQ2)** Do all proposed parameterized graph augmentation schemes benefit the contrastive learning process? How does each one affect the model performance?
- **(RQ3)** Can the proposed parameterized graph augmentation schemes generate meaningful augmented graph views?
- **(RQ4)** Are the proposed constraints make explainable augmentations? How do they influence specific molecules?

### 4.1 Experimental setups

#### 4.1.1 Datasets

##### Contrastive learning dataset

To train our MolCLE model, a large number of unlabeled molecule data is necessary to perform contrastive learning. Thus, we follow Hu et al. [23] to use 2 million unlabeled molecules^1^ sampled from the ZINC15 database [50] for graph contrastive learning.

##### Fine-tuning tasks and datasets

To prove the effectiveness of our MolCloze model, we select six molecule property prediction tasks from the MoleculeNet benchmark^2^ [60]. We summarize the used benchmark datasets in Table 2.

**Table 2:** Dataset summary.

| Category | Dataset | #Tasks | #Molecules | Metrics |
| --- | --- | --- | --- | --- |
| Biophysics | BBBP | 1 | 2039 | ROC-AUC |
| Physiology | BACE | 1 | 1513 | ROC-AUC |
|  | Tox21 | 12 | 7831 |  |
|  | ToxCast | 617 | 8575 |  |
|  | ClinTox | 2 | 1478 |  |
|  | SIDER | 27 | 1427 |  |

##### Fine-tuning dataset splitting

Similar to implementation in Rong et al. [47], we also use scaffold-splitting [3] to avoid obtaining overly optimistic results, which clusters compounds by scaffold (molecular graph substructure) and reassemble the clusters by placing the most common scaffolds in the training datasets, generating validation and test datasets which include structurally different compounds. Previous research works have demonstrated that this special scaffold splitting is more approximate to the ideal chronological split, which can stimulate the real-world application scenarios better. The model evaluated in the setting of scaffold splitting is convinced to give more reliable predictions. Therefore, in our research work, we use the special scaffold splitting to split the downstream benchmark datasets. The split ratio for train, validation and test datasets are typically 80%, 10%, and 10% respectively.

#### 4.1.2 Implementation details

In our experiments, we employ GIN with the default setting in Xu et al. [64] as the GNN encoder [23, 69]. Experiments are performed 10 times with mean and standard deviation of ROC-AUC scores (%) reported as Hu et al. [23]. We finetuned our MolCLE model on the training datasets of downstream molecular property prediction tasks. We use a batch size of 32 and dropout 0.5. We use Adam optimizer [28] for both contrastive learning and fine-tuning. All variables are initialized with Xavier [14]. We use the validation loss to select the best model. All models are implemented using PyTorch Geometric [11] and PyTorch [41]. All fine-tuning datasets are available in PyTorch Geometric libraries. The initial learning rates of contrastive learning and fine-tuning are 0.0001 and 0.001, respectively. All experiments are implemented on 1 NVIDIA Titan RTX GPU.

#### 4.1.3 Baselines

We comprehensively evaluate MolCLE against 8 popular supervised counterparts from MoleculeNet [60] and 3 state-of-the-arts self-supervised baselines. Among them, TF_Robust [44] is a DNN-based multitask framework taking the molecular fingerprints as the input. GraphConv [29], Weave [27] and SchNet [48] are three graph convolutional models. MPNN [12] and its variants DMPNN [66] and MGCN [34] are models considering the edge features during message passing. AttentiveFP [63] is an extension of the graph attention network. Specifically, to demonstrate the superiority of our proposed contrastive learning framework and parameterized graph augmentation schemes, we also compare MolCLE with three self-supervised models: Pre-GIN [23], GraphCL [69] and GROVER^3^ [47].

### 4.2 (RQ1) Performance results compared to baselines on multiple datasets

The experimental results are summarized in Table 3. Overall, from the table, we can see that our proposed model achieves strong performance across all six datasets. Moreover, our MolCLE outperforms all the supervised counterparts by considerable margins on molecular property prediction tasks. Besides, compared with Pre-GIN [23], our MolCLE pushes the performance more forward, demonstrating the superiority of the proposed contrastive learning framework. Furthermore, our MolCLE model achieves higher results across all datasets compared to GraphCL [69], which can be attributed to our proposed parameterized graph augmentation schemes, verifying their significance on molecular graph contrastive learning. We will analyze these schemes in detail when answering the following research questions. Finally, we can notice that our MolCLE model shows a comparative performance to GROVER [47], which has the largest GNNs architecture tailored for molecular graph data with tens of millions of parameters. However, our MolCLE, less than one million parameters, outperforms GROVER over three datasets and achieves similar results across the others, performing a notable success on extending contrastive learning to molecular graph data.

**Table 3:**
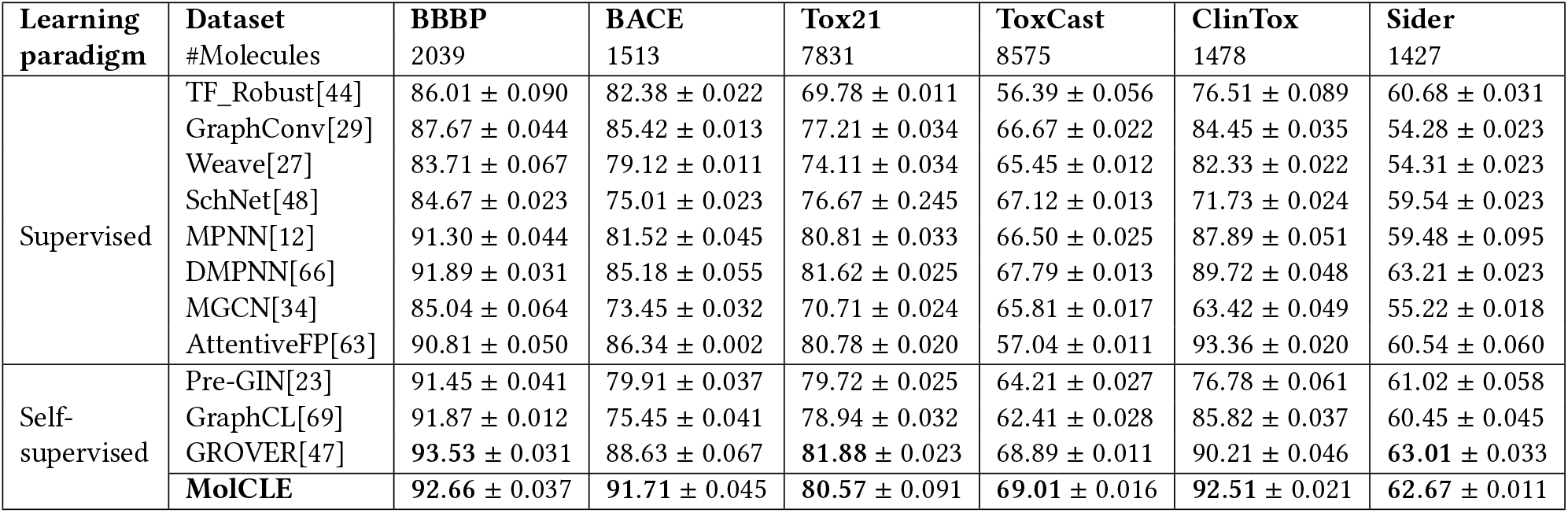
The performance results compared to baselines (ROC-AUC(%)).

### 4.3 (RQ2) Ablation studies on parameterized graph augmentation schemes

To answer this research question, we conduct ablation studies regarding our proposed parameterized graph augmentation schemes, where we substitute the parameterized topological and attributive augmentation with random edge perturbation and attribute masking. The experimental results are shown in Table 4. Obviously, our two parameterized augmentation components regarding topology and attribute all achieve performance improvements. Comparing *MolCLE-TA* with *MolCLE-AA*, we can conclude that the parameterized topological augmentation plays a more important role than the attributive one. The model variant without both these two parameterized schemes obtains the poorest performance which indicates the superiority of our proposed two augmentation schemes.

**Table 4:** Ablation studies on parameterized graph augmentation schemes (ROC-AUC(%)).

| Model variants | Topo. | Attr. | BBBP | BACE | Tox21 | ToxCast | ClinTox | Sider |
| --- | --- | --- | --- | --- | --- | --- | --- | --- |
| MolCLE-TA-AA | rand. | rand. | 91.25 ± 0.051 | 75.83 ± 0.035 | 78.81 ± 0.039 | 62.34 ± 0.026 | 85.26 ± 0.057 | 60.97 ± 0.044 |
| MolCLE-TA | rand. | Param. | 91.85 ± 0.012 | 76.48 ± 0.039 | 79.17 ± 0.044 | 64.53 ± 0.031 | 86.69 ± 0.024 | 61.39 ± 0.023 |
| MolCLE-AA | Param. | rand. | 92.43 ± 0.052 | 90.96 ± 0.048 | 79.89 ± 0.018 | 68.78 ± 0.020 | 91.95 ± 0.034 | 62.14 ± 0.031 |
| **MolCLE** | Param. | Param. | **92.66 ± 0.037** | **91.71 ± 0.045** | **80.57 ± 0.091** | **69.01 ± 0.016** | **92.51 ± 0.021** | **62.67 ± 0.011** |

### 4.4 (RQ3) Meaningful augmented graph views

Here we provide many molecule examples to illustrate what augmented graph views our MolCLE model generates. Figure 3 shows two augmented views for each of the three molecular graphs in a column. As shown in this figure, we can find that, although the two augmented views are very similar, they still remain slightly different, which demonstrates our proposed parameterized graph augmentation can generate effective positive pairs. Besides, the selected fragments for each molecular graph are obviously preserving chemical meanings. As shown in Figure 3, effective functional groups like benzene ring, carbonyl, hydroxy, carboxyl and halogen are all detected. On the opposite, futile fragments like carbon chains are ignored. Finally, we can notice that all of the augmented graph views keep a certain ratio with the original topological size, instead of select only a few bonds or all bonds.

**Figure 3:**
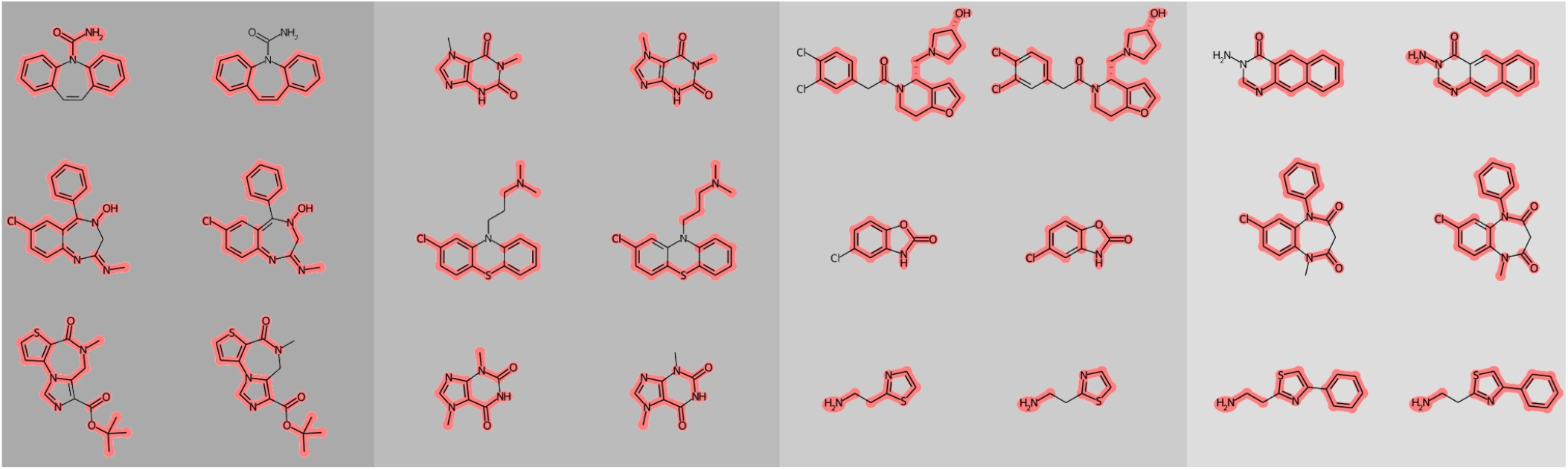
Generated views from two topological augmentation components in MolCLE. The two graph views from the same molecule serve as positive pairs during contrastive learning.

### 4.5 (RQ4) Explaining chemical intuitions from constrained augmentations

We demonstrate that MolCLE generates explainable augmentations with chemical meanings by case studies on the *BBBP* dataset. Various visualization examples verify the effectiveness of the proposed constraints to preserve chemical priors in generated augmentations automatically, as summarized in Figure 4.

**Figure 4:**
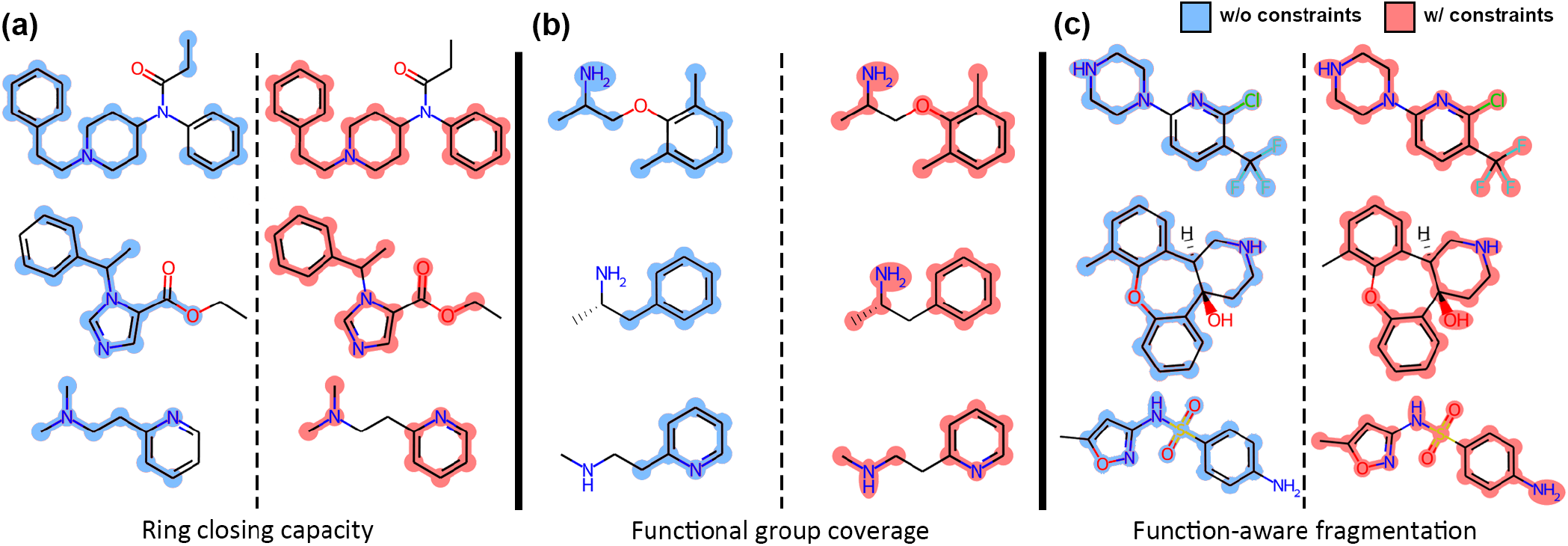
A comparison of topological augmentations learned from MolCLE without (blue) and with (red) the proposed constraints. With constraints, MolCLE (a)tends to generate augmentations with closed aromatic rings, such as phenyl, imidazolyl, and pyridineyl groups; (b)learns to cover all functional groups rather than the linkage atoms, e.g., selecting amine groups and aromatic rings on both sides of the molecule but not the alkyl groups between them; (c)decomposes a whole molecule into non-overlapping but property-preserving functional groups by bonds even without linkage atoms between functional groups.

We first concentrate on the most important aromatic rings in organic chemistry. On the left column of Figure 4, MolCLE learns to generate graph views preserving aromaticity, which allows contrastive training to learn the correct semantic information of the original molecule. Secondly, we demonstrate that, compared to generated graph views without constraints, MolCLE learns to cover the most meaningful functional groups in the molecule. As shown in the middle column in Figure 4, MolCLE could focus on both aromatic rings and the other amine group on the other side of the molecule. This result is in agreement with existing work that the property of the original molecule could be considered of the summation of the properties of its functional groups [9]. In addition, we find that molecule augmentations learned from MolCLE are decent molecular backbone fragments, similar to what the rule-based molecular fragmentation approach obtained [33]. This phenomenon reveals that our proposed parameterized graph augmentation scheme could decompose molecules from the connecting atoms, usually alkyl groups, to obtain a set of functional group fragments. As a matter of fact, molecular virtual fragmentation has proved to be an efficient way for *in silico* screening the chemical space in drug discovery, indirectly proving the effectiveness of our proposed parameterized graph augmentation scheme.

In conclusion, MolCLE could learn explainable molecule augmentations with chemical intuitions, providing a novel method to learn task-agnostic molecule embeddings from unlabeled data.

## 5 CONCLUSION

In this paper, we have developed a novel molecular graph contrastive learning framework with parameterized explainable augmentations. We pre-train our MolCLE model by maximizing the agreement between two augmented graph views from the same molecule. The proposed parameterized graph augmentation scheme self-adaptively learns topological and attributive augmentations for each molecule, which preserves the domain priors to make all augmented graph views chemically meaningful. The augmentation scheme can capture implicit graph connectivity patterns with chemical intuitions, and meanwhile, emphasize the underlying semantic information by selecting important feature dimensions. Our model outperforms many supervised counterparts and achieves a considerable performance compared to other state-of-the-art baselines. We also analyze the explainability of our generated graph views by case studies. Comprehensive experiments across multiple real-world datasets demonstrate the superiority of our model. In our future works, we will explore more contrastive learning frameworks and tailor a best-matched one for molecular graph data. Besides, we will try more downstream tasks like quantum chemical properties of molecules for better validate the generalizability and transferability of our proposed MolCLE model.

## Footnotes

1 The contrastive learning dataset is here: https://snap.stanford.edu/gnn-pretrain

2 All fine-tuning datasets can be found here: http://moleculenet.ai/datasets-16

3 We pre-train this model using the same dataset as our contrastive training stage.

## Notes

### Competing Interest Statement

The authors have declared no competing interest.

